# Solid Foods Drive the Maturation of the Gut Microbiota During Infancy

**DOI:** 10.64898/2026.09.09.750284

**Authors:** Yanan Wang, Heng Ku, Azadeh Safarchi, Megan Rebuli, Theodora Almond, Beverly Muhlhausler, Cuong D Tran, Michael Conlon

## Abstract

The establishment of a healthy gut microbiota in early life is an important determinant of long-term health, yet how the introduction of solid foods as a key nutritional transition shapes the gut microbiota development remains understudied. We aimed to characterize the impact of solid food introduction (complementary feeding) on the compositional succession and functional maturation of the infant gut microbiota during weaning. Infant faecal samples and corresponding dietary data were collected at five time points: pre-weaning (PREW, 4-6 months of age) 4, 8, and 12 weeks after first solid food introduction (WK4, WK8, WK12), and the late weaning stage (LATW, ∼12 months) when infants were transitioning to a family diet. Faecal microbiota composition was profiled using full-length 16S rRNA gene sequencing. Solid food introduction induced marked restructuring of the gut microbiota, characterized by significant increases in richness, evenness, and phylogenetic diversity (p < 0.01), together with the emergence of adult-associated taxa, including *Faecalibacterium*, *Ruminococcus*, and members of the Lachnospiraceae. These changes were most pronounced at late weaning and were strongly associated with increased intakes of plant-derived complex carbohydrates and dietary protein (Spearman correlation r > 0.30, FDR < 0.001). Butyrate-producing capability, a functional feature of gut microbiota maturation, was correlated with the establishment of key adult-like species, including *Ruminococcus bromii* and *Faecalibacterium prausnitzii* (Spearman correlation, r > 0.30, FDR < 0.001). Additionally, the reshaping of the gut microbiota by complementary feeding also attenuated microbial diversity associated with early-life factors, including delivery mode and household pet exposure. Together, these findings suggest that nutrient transitions during complementary feeding are key drivers of infant gut microbiota development and maturation, highlighting weaning as a critical window for shaping the gut microbiota and a potential target for early-life nutritional interventions to promote gut health and associated wellbeing.

## Introduction

The gut microbiota is closely associated with host health from birth [1] and plays important roles in the development of the immune system [2]. An unfavourable gut microbiota composition in early life has been linked to multiple childhood diseases and conditions including asthma [3, 4], allergic diseases [5, 6] and poor metabolic health [7, 8]. Therefore, establishing a healthy gut microbiota during early life may represent a promising strategy for disease prevention and the improved lifelong health outcomes [9–11].

The composition of the early-life microbiota is shaped by several perinatal and early-life factors. Mother’s gut microbiota, delivery mode, infant feeding practices (breast milk vs. formula feeding), skin/oral contact and the hospital/household environment collectively determine the initial microbial colonisation during early postnatal period [12, 13]. During this period, the gut microbiota is dominated by early colonisers including *Bifidobacterium*, *Bacteroides*, *Clostridium* and Enterobacteriaceae [12, 14]. Following the commencement of weaning, the period when an infant transitions from a solely milk-based diet to solid foods, the gut microbiota progressively develops toward an adult-like community, characterized by greater diversity and increased compositional and functional complexity [15]. Normal development of the gut microbiota during weaning underpins immune system maturation [16] and is critical to child health and long-term health outcomes [9].

Although the introduction of solid foods has been recognized as an important factor shaping the gut microbiota, how the associated nutrient transition shapes microbial succession and functional maturation remain understudied. One of the most important dietary changes during weaning is the introduction of plant-derived complex carbohydrates, such as dietary fibre. These dietary factors support the growth of adult-associated taxa, including *Ruminococcaceae* and *Lachnospiraceae* [17], and serve as fermentable substrates for the production of butyrate [17, 18], a short chain fatty acid (SCFA) that plays a crucial role in immune regulation and gut physiology [19–21]. The establishment of adult-associated taxa and increased butyrate-producing capacity are key features of gut microbiota maturation during this period [14, 22, 23]. Thus, understanding changes in key nutrients during weaning and their influence on gut microbiota succession and SCFA-producing capability is essential for elucidating how complementary feeding drives gut microbiota maturation and, in turn, identifying potential dietary approaches to optimise gut microbiota development.

Exclusive breastfeeding is recommended for the first six months of life [24]. However, in practice, solid foods are commonly introduced between 4 and 6 months [25, 26], resulting in considerable variation in weaning commencement timing among infants. Because gut microbial communities can respond rapidly to dietary change [27], thus investigating gut microbiota development according to weaning stage, rather than chronological age, provides a more biologically meaningful framework for understanding diet-microbiota interactions during this period.

In the current study, we therefore employed a weaning commencement-anchored longitudinal sampling strategy to characterize the developmental trajectory and ecological succession of the infant gut microbiota, focusing on key stages of weaning up to 12 months of age, when infants are transitioning to a family diet and the gut microbiota undergoes extensive development [15]. Combined with full-length PacBio 16S rRNA gene sequencing, this weaning stage-based approach enables high-resolution characterization of gut microbiota succession during weaning and addresses a critical knowledge gap regarding how nutrient transitions during this period drive early-life gut microbiota maturation.

## Methods

### Study population

Sixty-eight participants were screened and 32 healthy infants were recruited from Adelaide and surrounding areas (within 20 km of the Adelaide CBD), through social media (Facebook), local childcare and local family communities. Infants who were at least four months of age, had not commenced solid foods, were born full-term with normal birth weight (between 2,500 and 4,499 grams), and had no history of antibiotic use since birth were eligible to participate. Maternal antibiotic use during labour was recorded but not considered as an exclusion criterion. Ethical approval for this study was obtained from the Commonwealth Scientific and Industrial Research Organisation (CSIRO) Human Research Ethics Committee (2022_048_HREC). Written informed consent was obtained from a parent or legal guardian for each participant prior to study commencement.

### Demographic and birth event data collection

Following infant enrolment into the study, parents or legal guardians were provided with a questionnaire via REDCap to collect demographic data (e.g. sex, date of birth) and information on early-life factors previously shown to influence gut microbiota composition, including delivery mode, gestational age, birth weight, maternal antibiotic exposure during birth, household environment (presence of siblings and/or furry pets) and also sociodemographic information including family income and mother’s education level. Data regarding feeding practice were collected prior to PREW sample collection via a virtual interview.

### Faecal sample collection

Faecal samples and corresponding dietary data were collected at five time points: prior to the introduction of solid foods (pre-weaning, PREW); at 4, 8, and 12 weeks after the introduction of the first solid food (WK4, WK8, WK12); and at the late stage of weaning (LATW), when participating infants reached 12 months of age. Participating caregivers were provided with instructions for collecting faecal samples and asked to complete a questionnaire regarding infant’s and mother’s (if breasted at the time of sample collection) use of antibiotics and medication in the past four weeks prior to each sample collection. Faecal samples were collected by caregivers from their child’s nappy using a provided sterile container and spatula. Samples were then placed into a cooler bag with an ice pack and were collected by study staff within 24 hours. Following transfer to the analytical laboratory, samples were aliquoted for different analysis (e.g. DNA extraction, SCFA analysis) and stored at −80°C until further processing.

### Dietary assessment

Dietary data were collected using a weighed two-day food diary adapted from the OzFits study [28]. Caregivers were asked to record all solid foods, beverages, including breastmilk and formula, and supplements in a provided food diary booklet. Parents were asked to begin the food diary two days prior to the scheduled sample collection date, ensuring that dietary intake was recorded immediately preceding faecal sample collection. If no bowel movement occurred on the scheduled collection date, parents were asked to continue the food record until a faecal sample was obtained. Timeline for faecal sample collection and dietary assessment is shown in Fig. 1 (Icons of breastfeeding, diet and faecal samples are downloaded from Flaticon.com).

**Fig 1.**
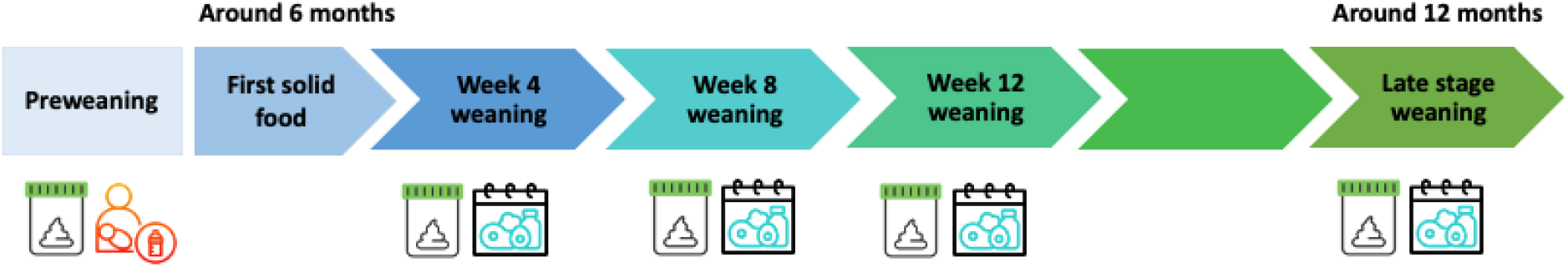
Timeline for faecal sample collection and dietary assessment.

Dietary data were entered into FoodWorks Professional (v10.0; Xyris Pty Ltd, Brisbane, Australia, 2019) by trained staff. The database used for dietary data processing was created based on several Data Sources in FoodWorks, including AusBrands 2019, AusFoods 2019, Abbott Products 2019, Nestle Baby Products 2019, Nutricia Advanced Medical Nutrition 2019 and Nutricia Early Life Nutrition 2019. For foods not available in the database, nutrient profiles of commercial products were entered manually based on nutritional information provided by the manufacturers; for homemade foods, individual ingredients and recipes were entered manually into the software to derive nutrient composition. Breastmilk intake was estimated using validated assumptions based on the duration of active suckling [28–30]. Briefly, parents were asked to record the duration of active suckling for each feed. Each feeding episode capped at a maximum of 10 min and feeds lasting less than 2 min were excluded from intake calculations. Feeds occurring within 30 min of each other were combined into a single feeding episode. Breastmilk intake was estimated assuming a milk transfer 10 mL/min with an energy density of 2.77 kJ/g [28]. Infant formula intake was calculated based on information provided by parents in the food diaries. Dietary data were subsequently checked for completeness, plausibility, and coding accuracy prior to analysis.

### Microbiome analysis

Faecal DNA was extracted using the QIAGEN PowerSoil kit (QIAGEN, Germantown, Maryland, USA) in accordance with the instructions provided by the manufacturer. The universal primer set, 27F (AGRGTTYGATYMTGGCTCAG) and 1492R (RGYTACCTTGTTACGACTT) was used to amplify the full-length 16S rRNA gene (V1–V9 regions) from genomic DNA [31], and the amplicon sequencing was performed using the Pacific Biosciences (PacBio) Sequel II long-read sequencing platform at the Australian Genome Research Facility (AGRF, Brisbane, Queensland, Australia). Raw PacBio HiFi reads were processed using the pb-16S-nf pipeline (version 0.6), incorporating QIIME 2-based analysis workflows [32]. Reads were quality filtered (quality score ≥20), and the number of reads per sample was limited to 50,000. Full-length 16S rRNA gene sequences were denoised using DADA2 [33], with a maximum expected error (maxEE) threshold of 10, to generate amplicon sequence variants (ASVs). Taxonomic classification of ASVs was performed using consensus alignment-based classification with VSEARCH against the Genome Taxonomy Database (GTDB R207), using a minimum sequence identity threshold of 97%.

### Faecal SCFA analysis

Faecal SCFA concentrations were determined using a method described by Watson et al. [34], with modifications. Briefly, approximately 250 mg of pre-weighed faecal sample was used for analysis. Heptanoic acid (1.68 mM) was used as the internal standard and added to the sample tubes at a volume equivalent to three times the weight of the faecal sample. Samples were vortex-mixed at high speed for 1 min and centrifuged at 2,000 × g for 10 min at 4 °C. The supernatants were collected and acidified with 10 µL of 1 M phosphoric acid. Subsequently, 300 µL of each sample was filtered using a Whatman PTFE 0.45 µm Mini-UniPrep filter. SCFA concentrations were quantified by gas chromatography (7890A; Agilent Technologies, Santa Clara, CA, USA) equipped with a flame ionisation detector and a Zebron ZB-FFAP capillary column (30 m × 0.53 mm × 1.00 µm; Phenomenex, Lane Cove, NSW, Australia).

### Statistical analysis

Group differences of beta-diversity of microbiota were assessed using the permutational analysis of variance (PERMANOVA) model with 9999 permutations based on the parameters permutation of residuals under a reduced model and a type III sum of squares (Primer-E v.7; Primer-E Ltd.). SIMPER (Similarity Percentages) was used to determine which taxa contributed most to differences between sample groups, with a cut-off set of cumulative contribution of 70% (Primer-E v.7; Primer-E Ltd.). Differences in alpha diversity and SCFA concentrations between each weaning stage (WK4, WK8, WK12, and LATW) and the PREW time point were assessed using the Kruskal-Wallis test, followed by Dunn’s post-hoc test for multiple comparisons. Group differences in alpha diversities between different life events (e.g., delivery mode, pet exposure), at each individual point was assessed using Mann-Whitney U test. A p-value < 0.05 was considered statistically significant. Spearman correlation analysis, followed by Benjamini-Hochberg false discovery rate (FDR) correction, with a threshold of 0.05, was used to assess correlations between nutrient intake and microbial species abundance, as well as between nutrient intake and microbiome alpha diversity.

## Results

### Characteristics of participating children

Infants participating in this study had an approximately equal sex distribution (53.1% male), and the majority were Caucasian (84.4%). All infants were born at full term (gestational age, 38.89 ± 0.91 weeks); 46.8% were born with Caesarean section, and 43.8% of infants were exclusively breastfed at enrolment prior to PREW sample collection (mean age, 4.61 ± 0.92 months). The mean age at introduction of solid foods was 5.60 ± 0.61 months, consistent with previous reports on the timing of complementary feeding among Australian infants [26]. The characteristics of the 32 participating infants are presented in Table 1.

**Table 1.** Characteristics of participating children (n=32).

|  |  |
| --- | --- |
| Sex-male, n (%) | 17 (53.1%) |
| Ethnicity- Caucasian, n (%) | 27 (84.4%) |
| Mother's education- Bachelor's degree or higher | 27 (84.4%) |
| Household income |  |
| *More than \$205, 000* | 7 (21.8%) |
| *\$105,001-\$205,000* | 20 (62.5%) |
| *\$70,001-\$105,000* | 5 (15.6%) |
| Delivery mode- Caesarean, n (%) | 15 (46.8%) |
| Gestational age (weeks), mean ± SD | $38.89 \pm 0.91$ |
| Birth weight (g), mean ± SD | $3406.47 \pm 446.50$ |
| Exclusive breastfeeding at enrolment, n (%) | 14 (43.8%) |
| Age at commencement of solids (months), mean ± SD | $5.60 \pm 0.61$ |

### Diversity and complexity of gut microbiota

A total of 156 out of 160 faecal samples with corresponding dietary data were collected across the five study time points. The mean age (months ± SD) of children at each collection point was as follows: PREW (4.93 ± 0.67), WK04 (6.60 ± 0.55), WK08 (7.55 ± 0.51), WK12 (8.45 ± 0.58), and LATW (12.32 ± 0.21). Two LATW samples and one WK12 sample were missing due to child illness, and one WK08 sample was missing due to family travel.

The faecal microbiota of participating children showed significant changes in diversity and community structure over the studied weaning period. Microbial communities at the five assessed weaning points were significantly different from each other based on unweighted UniFrac distances (Figure 2A, p =0.001), indicating continuous development of the gut microbiota following the commencement of solid foods. Alpha diversity increased progressively across the five assessed time points during weaning (Figure 2B-E) and all measured α-diversity indices were significantly higher at LATW compared to PREW (p <0.01). The gradual upward trend was more pronounced for metrics that are more sensitive to addition of new taxa, including Observed features and Phylogenetic diversity (PD), with significant increases detected from 8 weeks and 12 weeks post-introduction of solid foods, respectively (p <0.01). In contrast, increases in indices that take account of evenness (Pielou’s evenness and Shannon index) was significant at LATW (p <0.01 for Pielou’s evenness and p <0.0001 for Shannon).

**Fig 2.**
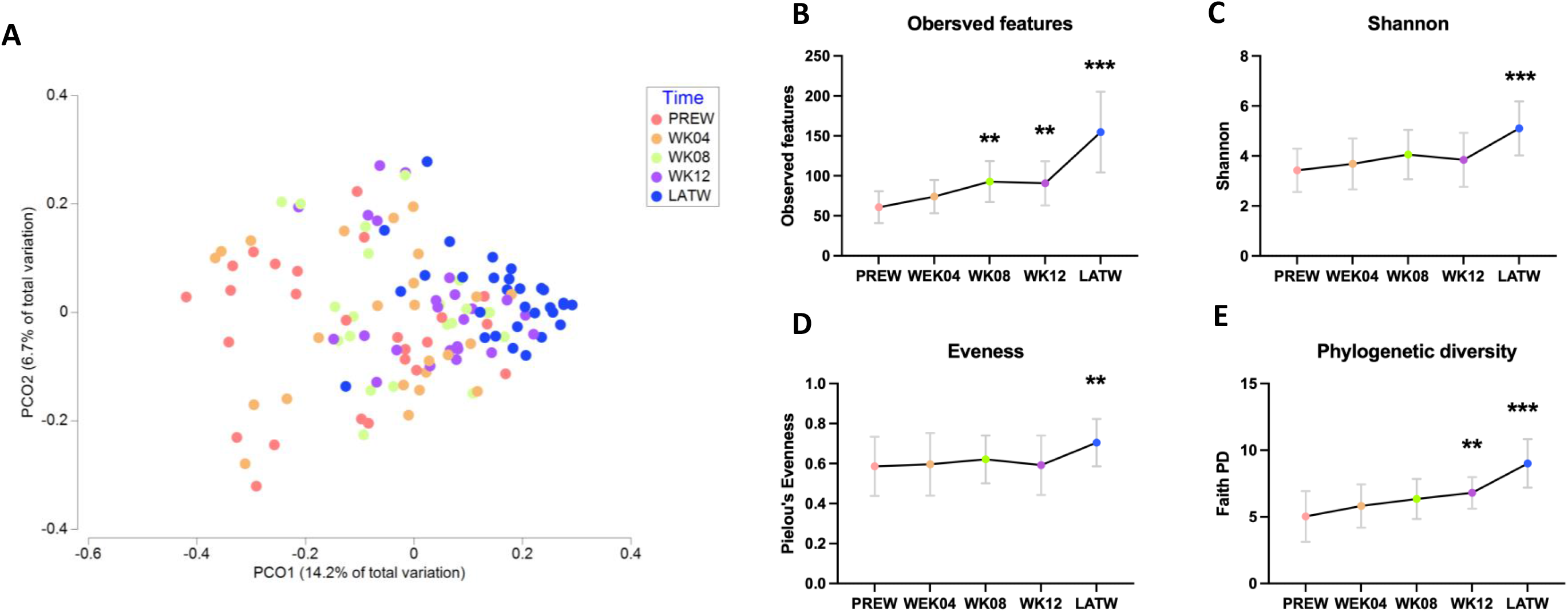
Changes in gut microbiota diversity during weaning. **A**. Beta diversity based on unweighted Unifrac distance. PERMANOVA, p =0.001. **B-E**, Alpha diversity indices, Kruskal-Wallis test, compared to PREW. **, p <0.01, ***, p <0.0001. PD, Phylogenetic diversity.

To capture temporal changes in taxa during the weaning period, we used SIMPER to identify genera that contributed to the separation of the microbial communities between two adjacent timepoints. Twenty-eight genera were identified as the primary contributors to community dissimilarities, based on a cumulative 70% contribution threshold (raw results of SIMPER were shown in Supplementary Table 1), and their relative abundances are shown in Figure 3.

**Fig. 3.**
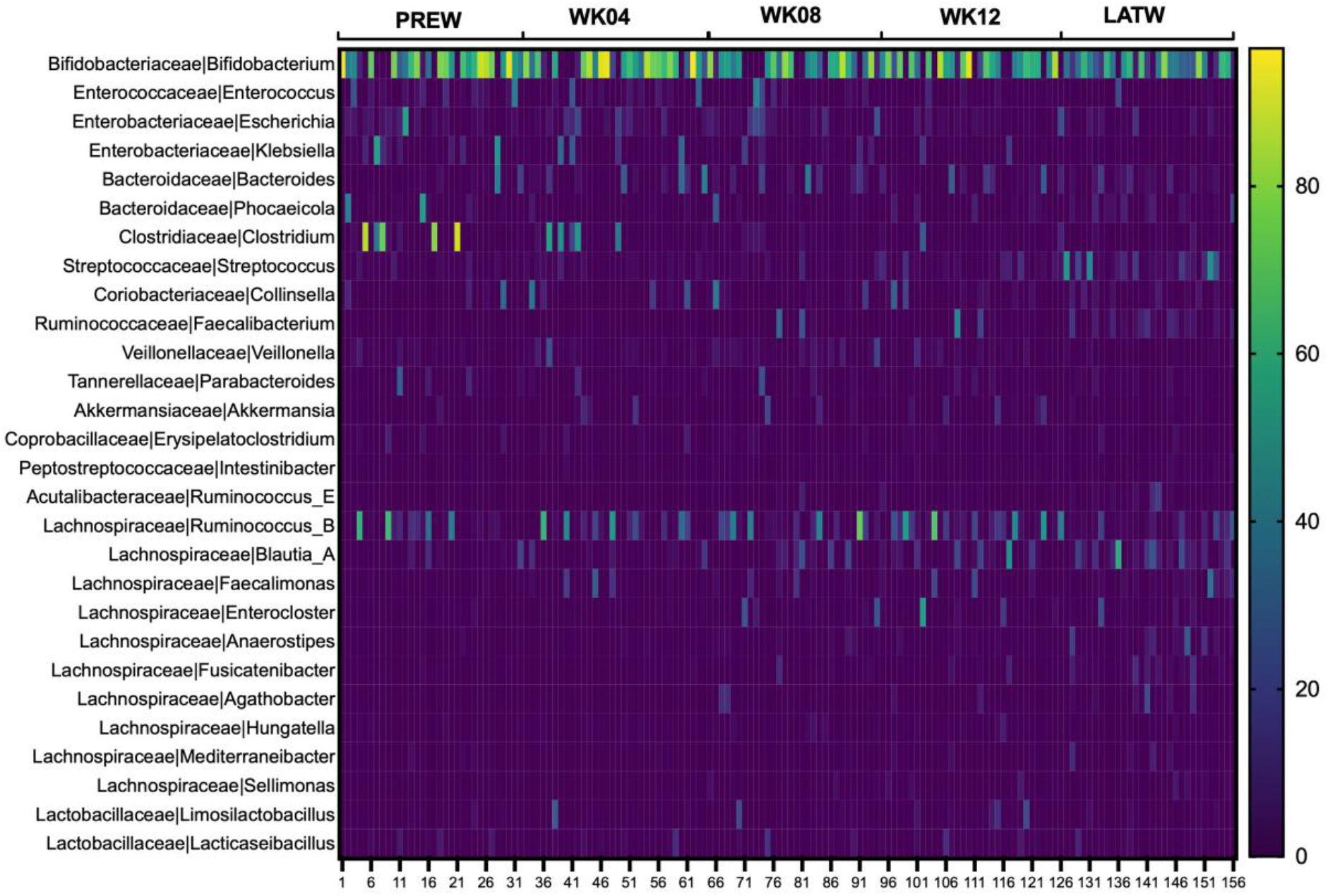
Relative abundance of taxa contributing to the dissimilarity between microbial communities at adjacent timepoints. Taxa were identified using SIMPER based on a 70% cumulative contribution threshold.

The infant faecal microbiota was dominated by *Bifidobacterium* throughout the weaning period, although its relative abundance declined markedly from PREW to LATW (median [IQR]: 50.72% [5.48–75.2%] at PREW vs 36.24% [13.77–60.88%] at LATW). Relative abundances of several early colonisers, including *Enterococcus*, *Escherichia*, *Klebsiella*, and *Phocaeicola*, declined over the weaning period. In contrast, taxa typically associated with a more mature, adult-like microbiota, such as *Blautia*, *Akkermansia*, and multiple members of the *Lachnospiraceae*, as well as *Faecalibacterium* and *Streptococcus*, increased progressively after the introduction of solid foods. Finally, *Bacteroides* and *Veillonella*, two commonly observed taxa in the infant gut microbiota, initially increased following the introduction of solids but subsequently declined by LATW.

### Changes in dietary intake and its association with the maturation of gut microbiota

Total energy, carbohydrate, and protein intake increased progressively across the weaning period, while total fat intake remained relatively stable (Fig. 4A). This resulted in a marked shift in macronutrient energy contribution from pre-weaning (PREW) to late weaning (LATW), characterized by a transition from fat to carbohydrates as the primary energy source and an increase in the proportional contribution of protein from 6.7% to 14.8% (Fig. 4B). The most pronounced dietary change during weaning was the introduction of complex carbohydrates, particularly starch and dietary fibre, which increased progressively across the weaning period from near-zero levels at baseline (Fig. 4C–D). In contrast, intakes of sugar remained unchanged across this same period (Fig. 4E). Dietary changes in food groups are shown in Supplementary Fig 1.

**Fig. 4.**
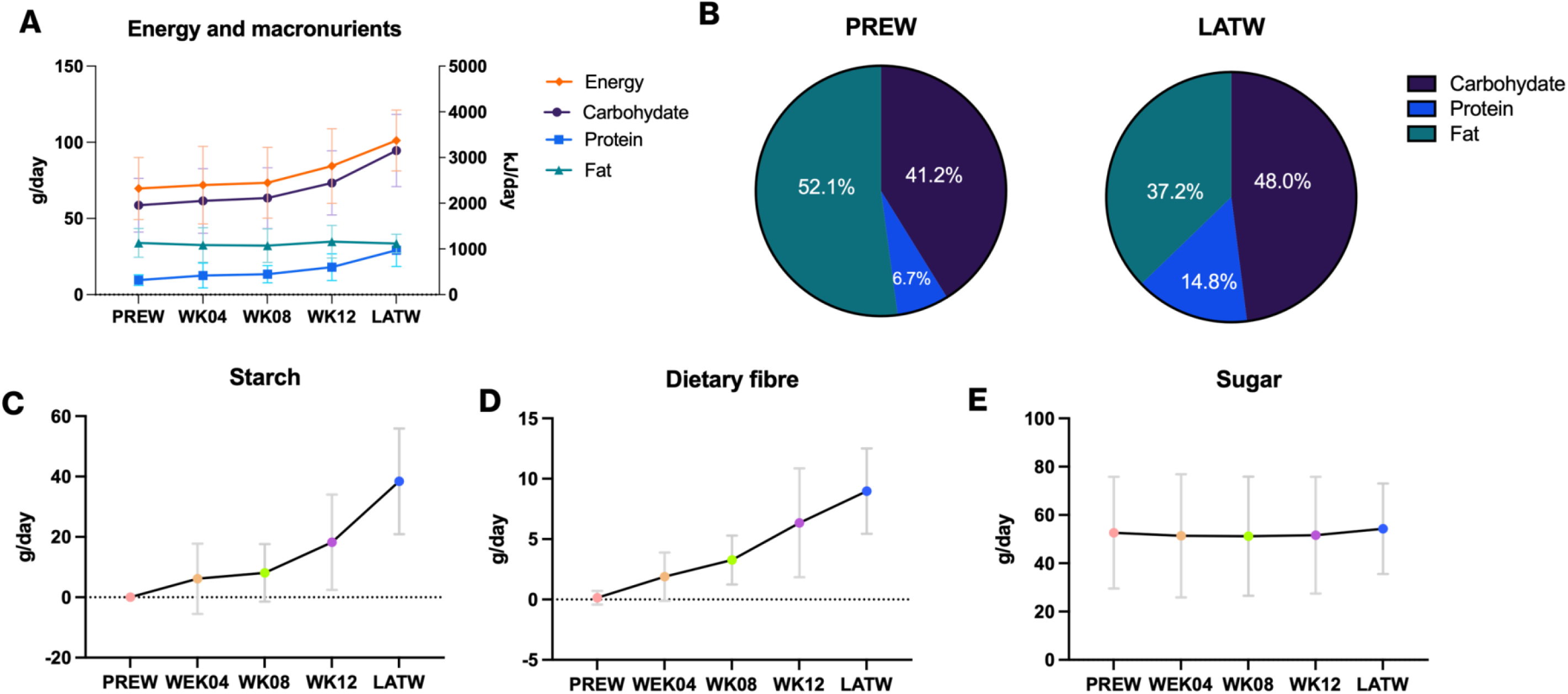
Changes in energy and macronutrient intake over the weaning period. (A) Energy and macronutrient intake over time. (B) Energy contribution from macronutrients at pre-weaning (PREW) and late weaning (LATW). (C–E) Changes in starch, dietary fibre, and sugar intake over time. Data are presented as mean ± SD.

The establishment of keystone species characteristic of a mature gut microbiota was strongly associated with the introduction of solid foods during weaning. This was reflected by moderate (r ≥0.30) to strong (r ≥0.50) positive correlations between specific taxa and the intake of newly introduced macronutrients, starch and dietary fibre, as well as protein, which increased significantly during this period (Fig 5). Notably, the associations with dietary fibre were particularly pronounced for *Ruminococcus bromii* (r =0.50, FDR <0.0001) and *Faecalibacterium prausnitzii* (r =0.48, FDR <0.0001), both recognized as key species involved in complex carbohydrate degradation and butyrate production. Typical adult-like taxa, including *Blautia wexlerae* and *Ruminococcus sp000433635*, also exhibited positive correlations with starch, dietary fibre, and protein intake (r ≥0.30, FDR <0.05). In addition to its association with complex carbohydrates, *Streptococcus thermophilus* demonstrated a strong positive correlation with protein intake (r = 0.58, FDR <0.0001). In contrast, species typically associated with breastfeeding, such as *Bacteroides fragilis* and *Phocaeicola dorei,* were negatively correlated with all three macronutrients (r < -0.30, FDR <0.05), suggesting that the changing microbial ecology during weaning no longer supports the persistence of these early-colonizing species.

**Fig. 5.**
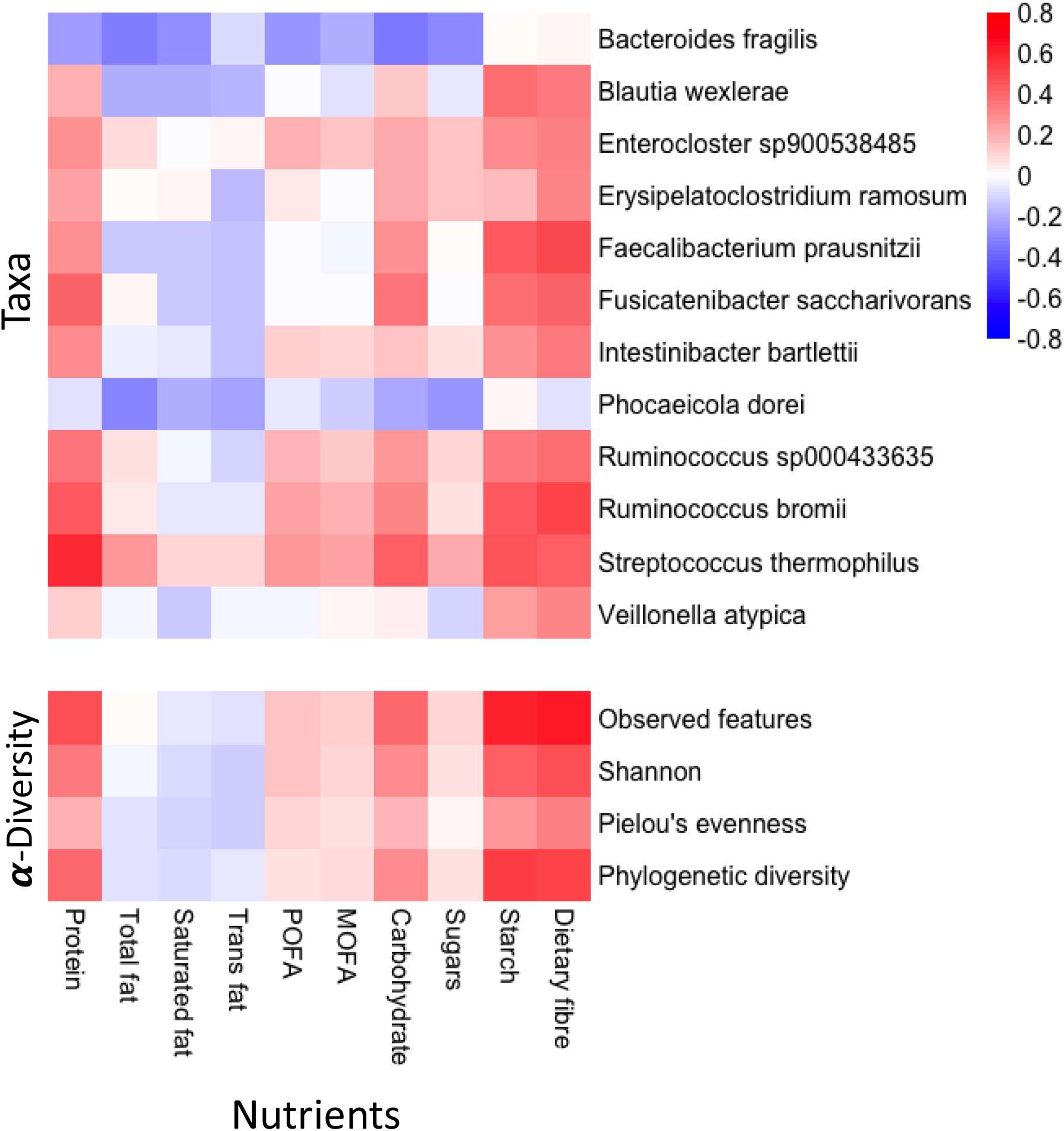
Spearman correlation between dietary macronutrients and gut microbiota. Upper panel: correlation with bacterial taxa at species level. Taxa that showed significant correlation with at least one nutrient are displayed (FDR-corrected *p* < 0.05, |*r*| ≥ 0.3). Lower panel: correlation with alpha diversity indices. Colour intensity reflects the correlation coefficient, ranging from −0.8 (blue) to +0.8 (red). * Indicates significant correlation with FDR <0.05. Spearman correlation analysis of the two panels was performed separately.

Microbial diversity, particularly richness and phylogenetic diversity, showed a strong response to the increased intake of carbohydrates (Fig 5). Observed features (indicating richness) and Phylogenetic diversity (Faith’s PD) were strongly correlated with both starch and dietary fibre (r >0.50, FDR <0.0001), and the correlation with Shannon and evenness were moderate and weak, respectively. In addition to carbohydrates, protein intake also showed significant correlation with richness (Observed features, r =0.47, FDR <0.0001), Shannon (r =0.34, FDR =0.00072) and Phylogenetic diversity (r =0.30, FDR <0.0001) of the gut microbiota.

### Key taxa driving the butyrate production during weaning

We assessed faecal short-chain fatty acid (SCFA) concentrations as a marker of gut microbiota maturation. While acetate and propionate levels remained stable throughout the weaning period, faecal butyrate concentrations were higher at 12 weeks following the introduction of solids (WK12) and at the late weaning stage (LATW) compared to pre-weaning (PREW) (Supplementary Fig. 2). Several taxa that responded markedly to the introduction of solid foods, namely *Ruminococcus bromii*, *Faecalibacterium prausnitzii*, *Clostridium* and *Lachnospiraceae* family, showed positive correlations with faecal butyrate levels (Fig. 6, FDR <0.01)

**Fig. 6.**
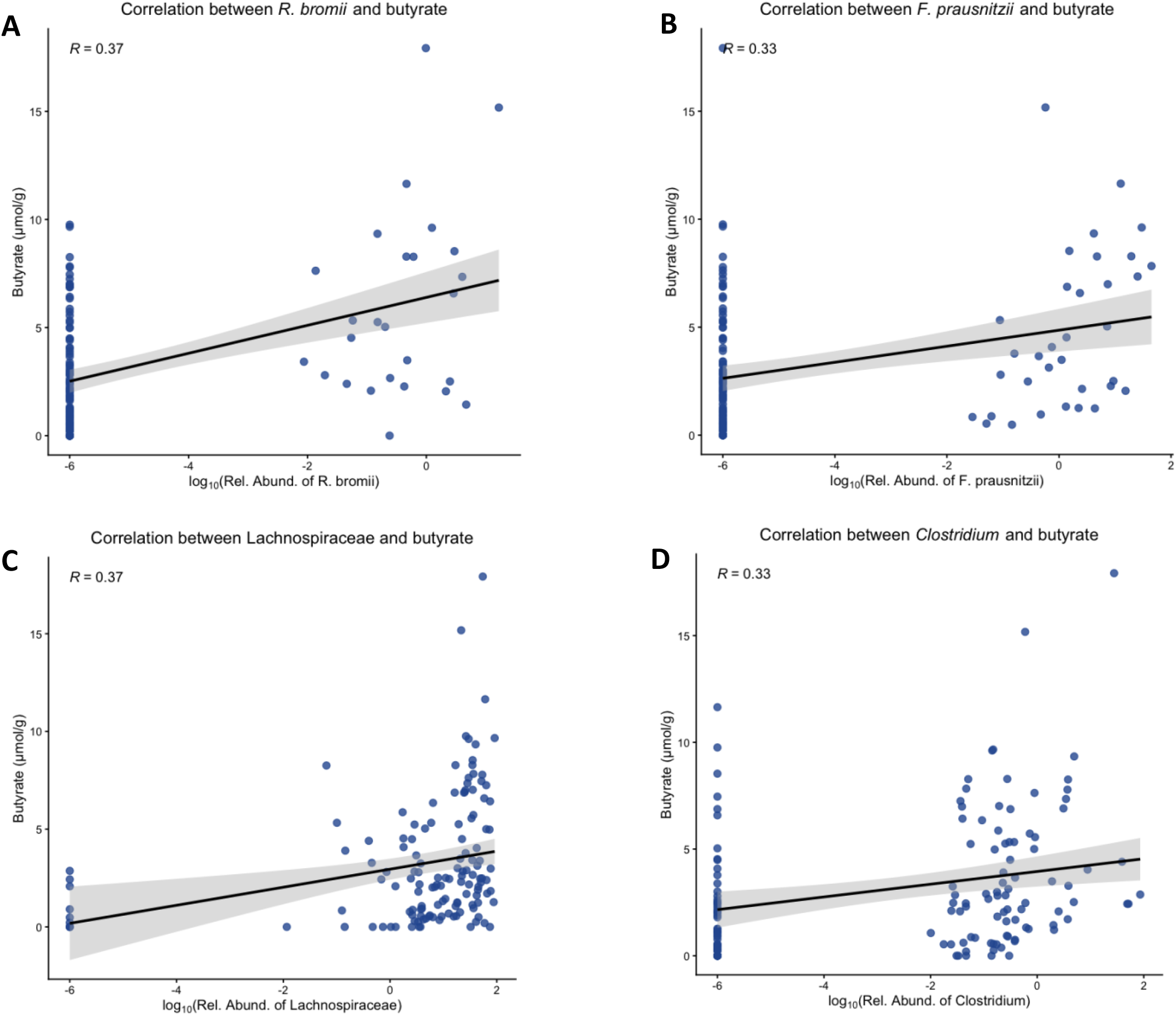
Potential butyrate-producing taxa during weaning and their correlations with faecal butyrate concentrations. Associations were assessed using Spearman correlation with false discovery rate (FDR) < 0.01

### Weaning diminishes distinct microbial diversity associated with delivery mode and furry pet exposure

Consistent with previous studies [35], we observed that infants born via vaginal delivery exhibited higher microbial diversity (Shannon) compared to those delivered by Caesarean section at 4 months of age before the introduction of solid foods. Similarly, infants living in households with furry pets showed greater phylogenetic diversity than those without at baseline (PREW). However, these alpha diversity indices were no longer different between groups within as little as 4 weeks following the introduction of solid foods (Fig. 7). However, feeding type and other early-life factors were not associated with microbiome diversity in this study.

**Fig. 7.**
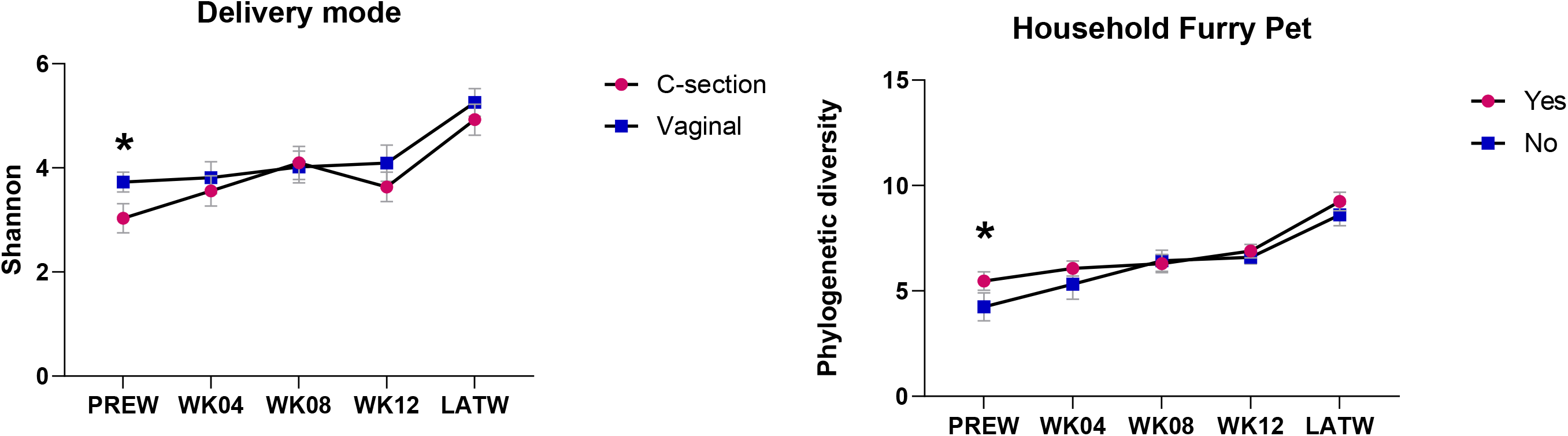
Differences in microbial diversity associated with delivery mode and household furry pet exposure diminished following the introduction of solid foods.

## Discussion

We characterized the detailed succession of gut microbiota during the weaning period (approximately between 6 and 12 months) when the gut microbiota undergoes its most extensive expansion in early life [15], and provided evidence of how changes in key nutrient during this period drive gut microbiota maturation.

Combined with longitudinal changes in faecal microbiota composition and detailed dietary records, we reported the dynamic ecological transition of the gut microbiota following the introduction of first solid foods. The ecological succession observed during weaning, characterised by a gradual decline in the relative dominance of *Bifidobacterium* and other early colonisers (e.g. *Enterococcus* and *Escherichia*), alongside increasing microbial diversity and expansion of taxa commonly associated with the adult gut microbiota, closely paralleled the major nutritional changes occurring during this period. The decline in *Bifidobacterium* was expected, as the progressive reduction in breast milk intake during weaning reduces the availability of human milk oligosaccharides (HMOs), an important substrate supporting the growth of this genus [36, 37]. At the same time, increasing consumption of plant-derived complex carbohydrates and dietary protein from complementary foods provides new ecological niches that favour the expansion of carbohydrate degraders and adult-like taxa, such as *Faecalibacterium*, *Ruminococcus*, and members of Lachnospiraceae. Importantly, key indicators of gut microbiota maturation, including increased microbial diversity and the emergence of the keystone butyrate-producing taxon *Faecalibacterium*, did not become evident until approximately eight weeks after the introduction of solid foods and were most pronounced at LATW, when the intake of complex carbohydrates and protein had increased substantially. These findings suggest that the ecological succession of the infant gut microbiota is closely linked to the progressive changes in nutrient availability during weaning. Rather than occurring immediately following the introduction of complementary foods, microbiota development appears to be a gradual process that unfolds over several weeks as infants establish a more diverse and complex diet.

Dietary fibre intake increased nearly linearly from the PREW to LATW (Fig. 4D) and was positively associated with both alpha diversity and the relative abundance of multiple adult-associated taxa (Fig. 5), supporting a central role for dietary fibre in promoting microbial diversification and the development of a more mature gut microbial community. At present, there are no established dietary fibre intake recommendations for infants under 12 months of age [38]. However, given the important roles of dietary fibre in shaping a healthy gut microbiota in early life and its potential benefits for children’s later life health, this area warrants greater attention and further investigation.

Starch intake exhibited associations with gut microbiota development that were broadly similar to those observed for dietary fibre, although the relationships were generally weaker. Grains and vegetables, particularly starchy and red or orange vegetables, were the most consumed complementary food during the weaning period (Supplementary Fig.1) and the main sources of starch. Although infants at this age possess a relatively high capacity to digest and absorb starch [5, 39, 40], approximately 5-20% of undigested starch can still reach to the large intestine [39, 41] and fuel the gut microbiota [42–44]. Similar to dietary fibre, starch appeared to serve as a driving force in promoting the increase in microbial diversity and increased abundance of multiple carbohydrate fermenter belonging to *Clostridium* cluster IV, including *F. prausnitzii* and *Ruminococcus spp.* Collectively, these findings suggest that the introduction of plant-derived complex carbohydrates during weaning is strongly associated with gut microbiota maturation, with increasing intake coinciding with the transition from a *Bifidobacterium*-dominated infant microbiota towards a more diverse community enriched in adult-associated, carbohydrate-degrading taxa.

The impact of dietary protein on gut microbiota development remains less well studied, particularly in young children. Unlike starch, the digestion and absorption of dietary protein in infants is less efficient than in adults, especially for proteins derived from complementary foods, due to the immaturity of the gastrointestinal digestive system [45, 46]. In our study, protein intake was positively associated with alpha diversity and the abundance of several members of the *Clostridia*, including *F. prausnitzii* and *Ruminococcus spp.*, which is broadly consistent with findings reported in Danish infants during the weaning period, where protein intake was also positively associated with microbiome diversity and the abundance of *Lachnospiraceae*, *Clostridiaceae*, and *Ruminococcaceae* [47]. Interestingly, in addition to the associations observed with the *Clostridia* taxa described above, we observed a strong positive correlation between protein intake and the relative abundance of *Streptococcus thermophilus*. The relative abundance of *S. thermophilus* was strongly associated with yogurt consumption (Suppl. Table 2), highlighting the potential contribution of dairy and fermented foods to the establishment of specific microbial taxa during weaning. Overall, although protein is not a preferred substrate for many taxa expanding during weaning, consistent with previous studies [47, 48], our findings suggest that increased consumption of protein-rich complementary foods, including meat and dairy products, may contribute to gut microbiota maturation alongside complex carbohydrates [49].

One important functional marker of gut microbiota maturation is an increased capacity for butyrate production [22, 50, 51]. Faecal butyrate levels of the infant participants in this study began to rise significantly from WK12 following the introduction of solid food and continued to rise through LATW. This is in line with the presence of primary butyrate producers (e.g., *Faecalibacterium*, *Ruminococcus*, and members of *Lachnospiraceae*)[52, 53] as well as the substantial increase in intake of complex carbohydrates and dietary fibre, which serve as the main substrates for butyrate production [52, 54]. The correlation test further confirmed the key taxa associated with butyrate production, including *R. bromii*, *F. prausnitzii*, Lachnospiraceae, and *Clostridium*, and their succession was significantly influenced by the introduction of solids. Butyrate production and high abundance of butyrate-producing bacteria have been consistently associated with a lower risk of immune-related diseases in later childhood, such as asthma and allergic diseases [55–57] and normal early-life development of the gut microbiota is thought to offer protective effects against these health problems later in life [50, 55, 58]. Our results highlight the importance of complementary feeding in driving the normal development of the gut microbiota and the potential for weaning to serve as a window of opportunity for the prevention of gut microbiota-associated diseases.

In addition, a notable finding of this study was that the influence of complementary feeding on gut microbiota development appeared sufficiently strong to diminish, and ultimately override, differences in microbial diversity associated with earlier life exposures, including delivery mode and household pet exposure. Both delivery mode and household pet exposure have previously been identified as important factors influencing early-life microbiota composition and diversity [35, 59, 60]. However, in the present study, their associations with gut microbiota diversity were no longer evident once complementary feeding was established, suggesting that the impact of these early-life exposures on microbial diversity do not necessarily persistent beyond early infancy, and has the potential to be modified or offset by influences beyond the immediate postnatal period, including complementary feeding.

This study has several limitations that should be considered. First, the relatively small sample size (n =32) limits statistical power, and further studies in larger populations are needed to confirm these findings. Second, the observational design limits our ability to draw conclusions about causal relationships. Additionally, dietary intake was assessed using a two-day food diary to reduce participant burden, and the relatively short recording period may compromise the accuracy of the dietary assessment. Nevertheless, the dietary patterns and nutritional changes observed in our study were highly consistent with those reported in previous studies using validated dietary assessment methods, supporting the reliability of our dietary findings.

Overall, our results highlight the potentially critical role of key nutrients introduced during complementary feeding in supporting the development and maturation of the gut microbiota. Normal gut microbiota maturation, marked by increased diversity, presence of keystone species such as *F. prausnitzii* and *R. bromii* and their production of the beneficial microbial metabolite butyrate, has been associated with a range of positive health outcomes. The establishment of these health-associated microbiome characteristics can be strongly influenced by complementary feeding. Therefore, complementary feeding during weaning warrants greater attention as a modifiable factor for establishing a healthy gut microbiota in early life. Our findings warrant further investigation of nutrient-microbe interactions during weaning to develop effective nutritional strategies for shaping a healthy gut microbiota and supporting host health during infancy and throughout life.

## Acknowledgements

We would like to thank Mr Himanshu Tandon and Dr Katie Wood for their assistance with participant recruitment and data management; Mr Tony Vu, Ms Cathryn Pape, and Ms Eliza Borgese for their assistance with faecal sample and dietary data collection; Ms Mariam for her laboratory support; and Dr Jatinder Sidhu and Dr Gupta Vadakattu for their scientific advice and support to the project. We would also like to thank the SAHMRI OzFITS Study team for kindly sharing their food diary protocol with our study.

## Supplementary Materials

**Suppl. Table 1**. SIMPER result output based on a cumulative 70% contribution threshold.

**Suppl. Table 2.** Spearman correlation between microbial species and food groups.

**Suppl. Fig 1.** Dietary changes in main food groups and subgroups.

**Suppl. Fig 2.** Changes in faecal SCFA concentrations during weaning.

## References

1. Arrieta, M.C., et al., The intestinal microbiome in early life: health and disease. Front Immunol, 2014. 5: p. 427.

2. Martin, R., et al., Early life: gut microbiota and immune development in infancy. Benef Microbes, 2010. 1(4): p. 367–82.

3. Stokholm, J., et al., Maturation of the gut microbiome and risk of asthma in childhood. Nat Commun, 2018. 9(1): p. 141.

4. Boulund, U., et al., The role of the early-life gut microbiome in childhood asthma. Gut Microbes, 2025. 17(1): p. 2457489.

5. Akagawa, S. and K. Kaneko, Gut microbiota and allergic diseases in children. Allergol Int, 2022. 71(3): p. 301–309.

6. Ke, H., H. Yao, and P. Wei, Advances in research on gut microbiota and allergic diseases in children. Curr Res Microb Sci, 2025. 8: p. 100362.

7. Jian, C., et al., Early-life gut microbiota and its connection to metabolic health in children: Perspective on ecological drivers and need for quantitative approach. EBioMedicine, 2021. 69: p. 103475.

8. Nunez, H., et al., Early life gut microbiome and its impact on childhood health and chronic conditions. Gut Microbes, 2025. 17(1): p. 2463567.

9. Borrego-Ruiz, A. and J.J. Borrego, Early-life gut microbiome development and its potential long-term impact on health outcomes. Microbiome Res Rep, 2025. 4(2): p. 20.

10. Yao, Y., et al., The Role of Microbiota in Infant Health: From Early Life to Adulthood. Front Immunol, 2021. 12: p. 708472.

11. Kerr, C.A., et al., Early life events influence whole-of-life metabolic health via gut microflora and gut permeability. Crit Rev Microbiol, 2015. 41(3): p. 326–40.

12. Enav, H., F. Backhed, and R.E. Ley, The developing infant gut microbiome: A strain-level view. Cell Host Microbe, 2022. 30(5): p. 627–638.

13. Martino, C., et al., Microbiota succession throughout life from the cradle to the grave. Nat Rev Microbiol, 2022. 20(12): p. 707–720.

14. Backhed, F., et al., Dynamics and Stabilization of the Human Gut Microbiome during the First Year of Life. Cell Host Microbe, 2015. 17(5): p. 690–703.

15. Ding, M., et al., Infant gut microbiome reprogramming following introduction of solid foods (weaning). Gut Microbes, 2025. 17(1): p. 2571428.

16. Al Nabhani, Z., et al., A Weaning Reaction to Microbiota Is Required for Resistance to Immunopathologies in the Adult. Immunity, 2019. 50(5): p. 1276–1288 e5.

17. Flint, H.J., et al., Microbial degradation of complex carbohydrates in the gut. Gut Microbes, 2012. 3(4): p. 289–306.

18. Louis, P. and H.J. Flint, Diversity, metabolism and microbial ecology of butyrate-producing bacteria from the human large intestine. FEMS Microbiol Lett, 2009. 294(1): p. 1–8.

19. Canani, R.B., et al., Potential beneficial effects of butyrate in intestinal and extraintestinal diseases. World J Gastroenterol, 2011. 17(12): p. 1519–28.

20. Mukhopadhya, I. and P. Louis, Gut microbiota-derived short-chain fatty acids and their role in human health and disease. Nat Rev Microbiol, 2025. 23(10): p. 635–651.

21. Koh, A., et al., From Dietary Fiber to Host Physiology: Short-Chain Fatty Acids as Key Bacterial Metabolites. Cell, 2016. 165(6): p. 1332–1345.

22. Appert, O., et al., Initial butyrate producers during infant gut microbiota development are endospore formers. Environ Microbiol, 2020. 22(9): p. 3909–3921.

23. Koenig, J.E., et al., Succession of microbial consortia in the developing infant gut microbiome. Proc Natl Acad Sci U S A, 2011. 108 **Suppl 1**(Suppl 1): p. 4578-85.

24. Organization, W.H., WHO Guideline for complementary feeding of infants and young children 6–23 months of age. 2023. p. 95.

25. Welfare, A.I.o.H.a., 2010 Australian National Infant Feeding Survey: indicator results. 2011: Canberra.

26. Netting, M.J., et al., The Australian Feeding Infants and Toddler Study (OzFITS 2021): Breastfeeding and Early Feeding Practices. Nutrients, 2022. 14(1).

27. David, L.A., et al., Diet rapidly and reproducibly alters the human gut microbiome. Nature, 2014. 505(7484): p. 559–63.

28. Moumin, N.A., et al., Usual Nutrient Intake Distribution and Prevalence of Inadequacy among Australian Children 0-24 Months: Findings from the Australian Feeding Infants and Toddlers Study (OzFITS) 2021. Nutrients, 2022. 14(7).

29. Thomas Berube, L., et al., Concerns About Current Breast Milk Intake Measurement for Population-Based Studies. J Acad Nutr Diet, 2018. 118(10): p. 1827–1831.

30. Byrne, R., A. Magarey, and L. Daniels, Food and beverage intake in Australian children aged 12-16 months participating in the NOURISH and SAIDI studies. Aust N Z J Public Health, 2014. 38(4): p. 326–31.

31. Buetas, E., et al., Full-length 16S rRNA gene sequencing by PacBio improves taxonomic resolution in human microbiome samples. BMC Genomics, 2024. 25(1): p. 310.

32. Pacific Biosciences. pb-16S-nf: Nextflow pipeline for PacBio HiFi full-length 16S rRNA gene analysis. 2022; 0.6:[Available from: https://github.com/PacificBiosciences/pb-16S-nf.

33. Callahan, B.J., et al., DADA2: High-resolution sample inference from Illumina amplicon data. Nat Methods, 2016. 13(7): p. 581–3.

34. Watson, E.J., et al., Human faecal collection methods demonstrate a bias in microbiome composition by cell wall structure. Sci Rep, 2019. 9(1): p. 16831.

35. Azad, M.B., et al., Gut microbiota of healthy Canadian infants: profiles by mode of delivery and infant diet at 4 months. CMAJ, 2013. 185(5): p. 385–94.

36. Sela, D.A. and D.A. Mills, Nursing our microbiota: molecular linkages between bifidobacteria and milk oligosaccharides. Trends Microbiol, 2010. 18(7): p. 298–307.

37. Lordan, C., et al., Linking human milk oligosaccharide metabolism and early life gut microbiota: bifidobacteria and beyond. Microbiol Mol Biol Rev, 2024. 88(1): p. e0009423.

38. National Health and Medical Research Council. Dietary fibre. Eat for Health [cited 2026 1 July]; Available from: https://www.eatforhealth.gov.au/nutrient-reference-values/nutrients/dietary-fibre.

39. Shulman, R.J., Starch Malabsorption in Infants. J Pediatr Gastroenterol Nutr, 2018. 66 **Suppl 3**: p. S65–S67.

40. Lin, A.H.-M. and B.L. Nichols, The digestion of complementary feeding starches in the young child. Starch - Stärke, 2017. 69(7-8): p. 1700012.

41. Christian, M.T., et al., Modeling 13C breath curves to determine site and extent of starch digestion and fermentation in infants. J Pediatr Gastroenterol Nutr, 2002. 34(2): p. 158–64.

42. Liu, S., et al., Starch and starch hydrolysates are favorable carbon sources for bifidobacteria in the human gut. BMC Microbiol, 2015. 15: p. 54.

43. Wang, Y., et al., The Capacity of the Fecal Microbiota From Malawian Infants to Ferment Resistant Starch. Front Microbiol, 2019. 10: p. 1459.

44. Christian, M.T., et al., Starch fermentation by faecal bacteria of infants, toddlers and adults: importance for energy salvage. Eur J Clin Nutr, 2003. 57(11): p. 1486–91.

45. Lee, S., et al., Low Protein Digestibility of Beef Puree in Infant In Vitro Digestion Model. Food Sci Anim Resour, 2019. 39(6): p. 1000–1007.

46. Asensio-Grau, A., et al., Complementary feeding in infants with cystic fibrosis: In vitro nutrient digestibility and impact on colonic microbiota. Food Bioscience, 2024. 59: p. 104249.

47. Laursen, M.F., et al., Infant Gut Microbiota Development Is Driven by Transition to Family Foods Independent of Maternal Obesity. mSphere, 2016. 1(1).

48. Clarke, S.F., et al., Exercise and associated dietary extremes impact on gut microbial diversity. Gut, 2014. 63(12): p. 1913–20.

49. Jackson, R., et al., Protein combined with certain dietary fibers increases butyrate production in gut microbiota fermentation. Food Funct, 2024. 15(6): p. 3186–3198.

50. Ruohtula, T., et al., Maturation of Gut Microbiota and Circulating Regulatory T Cells and Development of IgE Sensitization in Early Life. Front Immunol, 2019. 10: p. 2494.

51. Nilsen, M., et al., Butyrate Levels in the Transition from an Infant- to an Adult-Like Gut Microbiota Correlate with Bacterial Networks Associated with Eubacterium Rectale and Ruminococcus Gnavus. Genes (Basel), 2020. 11(11).

52. Louis, P. and H.J. Flint, Formation of propionate and butyrate by the human colonic microbiota. Environ Microbiol, 2017. 19(1): p. 29–41.

53. Singh, V., et al., Butyrate producers, “The Sentinel of Gut”: Their intestinal significance with and beyond butyrate, and prospective use as microbial therapeutics. Front Microbiol, 2022. 13: p. 1103836.

54. Zhang, M., et al., Recent advances in developing butyrogenic functional foods to promote gut health. Crit Rev Food Sci Nutr, 2024. 64(13): p. 4410–4431.

55. Depner, M., et al., Maturation of the gut microbiome during the first year of life contributes to the protective farm effect on childhood asthma. Nat Med, 2020. 26(11): p. 1766–1775.

56. Bankole, T. and Y. Li, The early-life gut microbiome in common pediatric diseases: roles and therapeutic implications. Front Nutr, 2025. 12: p. 1597206.

57. Arrieta, M.C., et al., Early infancy microbial and metabolic alterations affect risk of childhood asthma. Sci Transl Med, 2015. 7(307): p. 307ra152.

58. Dogra, S.K., et al., Nurturing the Early Life Gut Microbiome and Immune Maturation for Long Term Health. Microorganisms, 2021. 9(10).

59. Stewart, C.J., et al., Temporal development of the gut microbiome in early childhood from the TEDDY study. Nature, 2018. 562(7728): p. 583-588.

60. Azad, M.B., et al., Infant gut microbiota and the hygiene hypothesis of allergic disease: impact of household pets and siblings on microbiota composition and diversity. Allergy Asthma Clin Immunol, 2013. 9(1): p. 15.

