## Supplementary figures and images for "Solid Foods Drive the Maturation of the Gut Microbiota During Infancy"

### Suppl Fig 1

#### Slide 1
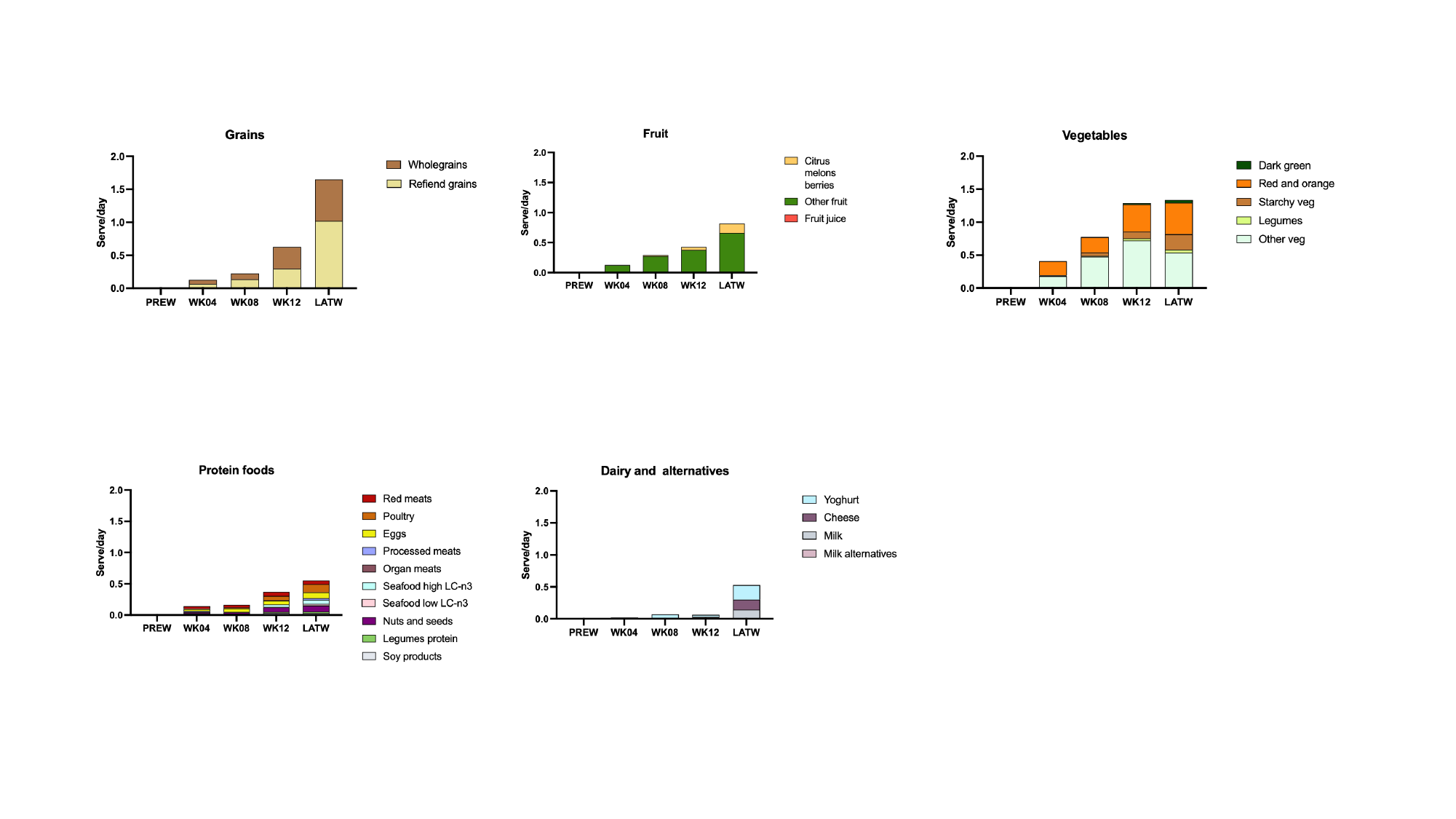

### Suppl Fig 2

#### Slide 1
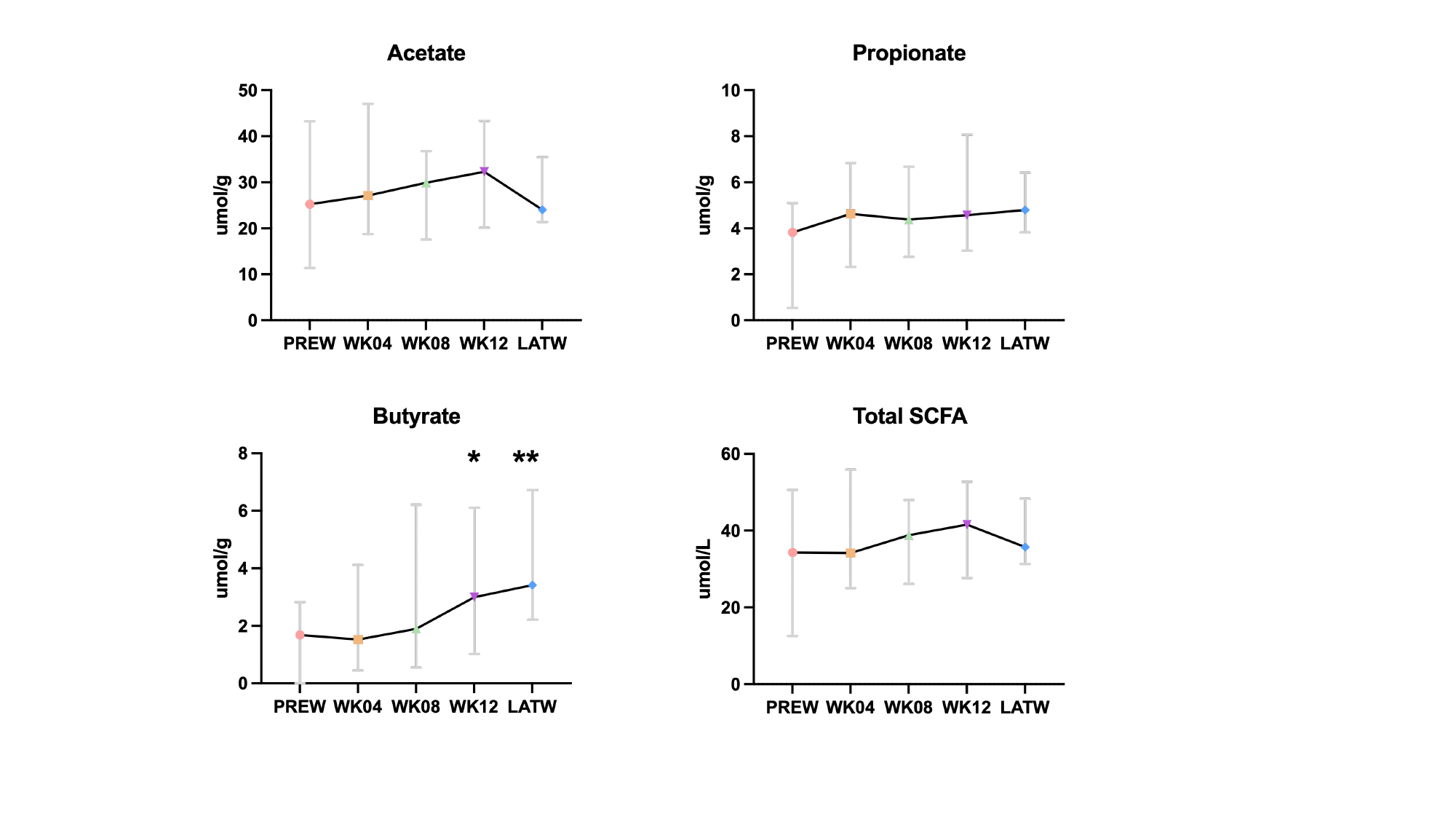
